# Regulatory vs. helper CD4^+^ T-cell ratios and the progression of HIV/AIDS disease

**DOI:** 10.1101/006676

**Authors:** Wilfred Ndifon, Jonathan Dushoff, Daniel Coombs

**Author notes:** **Address for correspondence:** Wilfred Ndifon, African Institute for Mathematical Sciences, 5-7 Melrose Road, Muizenberg 7945, Cape Town, South Africa.

## Abstract

The causes of individual variability in the length of time between human immunodeficiency virus (HIV) infection and the development of AIDS are incompletely understood. Here, we present a novel hypothesis: that the relative magnitude of responses to HIV mediated by CD4^+^ T regulatory (Treg) cells vs. CD4^+^ T effector (Teff) cells is a critical determinant of variability in AIDS progression rates. We use a simple mathematical model to show that this hypothesis can plausibly explain three qualitatively different outcomes of HIV infection – fast or slow progression to AIDS, and long-term non-progression to AIDS – based on individual variation in underlying T-cell response. This hypothesis also provides a unifying explanation for various other empirical observations, suggesting in particular that both aging and certain dual infections increase the rate of AIDS progression because they increase the strength of the Treg cell response. We discuss potential therapeutic implications of our results.

## Introduction

The length of time that it takes for different untreated patients infected with human immunodeficiency virus (HIV) to develop AIDS varies from <2 y to >20 y, with a median of about 10 y (CASCADE 2000; Murray et al. 2005). Progression to AIDS in individual patients is generally characterized by the occurrence of specific opportunistic infections coincident with a sharp rise in the amount of HIV found in the blood (the HIV load) and a precipitous decline in the frequencies of CD4^+^ T cells (Murray et al. 2005; Grabar et al. 2009). A small fraction of HIV patients – called elite HIV controllers – do not develop AIDS for a very long time after being infected, and they also maintain undetectable levels of HIV in their blood as well as normal frequencies of CD4^+^ T cells (Grabar et al. 2009). Studies conducted during the past 30 y have identified a variety of virus and host factors that contribute to this individual variability in AIDS progression rates (Cornelissen et al. 2012; Hogan & Hammer 2001; Jaffar et al. 2004; Kosmrlj et al. 2010; Mellors et al. 1997; Migueles et al. 2000; Migueles et al. 2008; Samson et al. 1996; Toossi et al. 2001; Vasan et al. 2006).

A number of studies have revealed associations between HIV viral characteristics and the rate of AIDS progression. For example, HIV type 1, the most prevalent type, tends to cause disease faster than type 2, which occurs mostly in sub-Saharan Africa (Jaffar et al. 2004), while among type 1 viruses, those of subtype D tend to cause disease faster than other subtypes (Vasan et al. 2006). Furthermore, it has been shown that after being infected by certain HIV variants, individuals that are subsequently co-infected by certain other pathogens, including *Mycobacterium tuberculosis* and other HIV variants, tend to progress to AIDS faster compared to individuals without such co-infections (Cornelissen et al. 2012; Toossi et al. 2001).

Other studies have implicated host genetic co-factors in AIDS progression. For example, certain mutations in a host’s CCR-5 chemokine co-receptor gene, which encodes one of the receptors used by HIV to infect target cells, are associated with a slower rate of development of AIDS (Samson et al. 1996). Another relevant factor is the dominant HLA I allele used by HIV-specific CD8^+^ T cells: hosts whose HIV-specific CD8^+^ T cells are restricted to the HLA-B57 allele tend to develop AIDS much more slowly than other patients (Hogan & Hammer 2001; Migueles et al. 2000), possibly because the HLA-B57-restricted cells recognize a larger diversity of HIV mutants (Kosmrlj et al. 2010).

Characteristics of the host’s immune response to HIV have also been associated with variability in the rate of AIDS progression. HIV-specific CD8^+^ T cells prevent HIV replication *in vitro* by lysing infected cells (Walker et al. 1986), and CD8^+^ T cells from AIDS nonprogressors are more effective at lysing HIV-infected cells, through the release of perforin and granzyme B, compared to CD8^+^ T cells from progressors (Migueles et al. 2008). CD4+ T cells have also been implicated in AIDS progression, with CD4^+^ T cell counts being strongly correlated with AIDS progression (Mellors et al. 1997). However, it is unclear whether the decline in CD4^+^ T counts is a cause or an effect of AIDS progression, or both.

Several theoretical studies have examined various aspects of the interactions between CD4^+^ T cells and HIV that occur during HIV infection. For example, by making a mathematical model that accounted for HIV killing of CD4^+^ T cells, the elimination of HIV by adaptive immune responses mediated by CD4^+^ T cells, and HIV evolution, Nowak and co-workers (Nowak & May 1991; Nowak et al. 1990) argued that progression to AIDS occurs when the antigenic diversity of HIV exceeds a defined threshold value. The mechanistic basis for this “diversity threshold” theory is that the elimination of HIV by adaptive immune responses is HIV strain-specific, whereas the impairment of immune responses by HIV is not (Nowak & May 1991; Nowak et al. 1990). Perelson and co-workers (Perelson et al. 1993) showed that variability in the rate of decline of CD4^+^ T cell counts, which is correlated with variability in AIDS progression rates (Mellors et al. 1997), can arise even in a mathematical model that does not account for either the immune response or HIV evolution, but considers only the infection of CD4^+^ T cells by HIV. Together, these theoretical studies imply that the dynamics of the interaction between HIV and CD4^+^ T cells might play a crucial role in the development of AIDS in individual patients.

Here, by focusing on the role of CD4+ T cells in the elimination of HIV by the immune system, we present a hypothesis that we show, using mathematical modeling, can explain empirical patterns of AIDS progression, as well as various other related empirical observations. In particular, these patterns emerge generically from simple assumptions based directly on experimental observations that: 1) The immunological “help” provided to CD8 and B cells by CD4^+^ CD25^lo^ Foxp3^−^ T effector (Teff) cells is generally contingent on the recognition of similar antigens by both the helper and the helped immune cell (Lanzavecchia 1985; Shedlock & Shen 2003; Murphy et al. 2011), and 2) CD4^+^ CD25^hi^ Foxp3^+^ T regulatory (Treg) cells suppress Teff cells in a “bystander” manner, largely independent of the antigen specificities of the targeted cells (Shevach 2009; Thornton & Shevach 2000). We will begin by describing our hypothesis verbally, and then proceed to analyze it mathematically.

## Results

### Hypothesis to explain individual variability in HIV/AIDS progression

Our hypothesis rests on the following experimental observations:

1. Teff cells enhance the functions of other effector immune cells that control HIV replication – e.g. by promoting the maturation of memory B and CD8+ T cells (Lanzavecchia 1985; Murphy et al. 2011; Shedlock & Shen 2003),
2. Treg cells suppress effector immune responses – e.g. by competing for IL-2 and secreting cytokines, such as TGF-β, which dampen the activation of effector cells (Murphy et al. 2011; Shevach 2009; Thornton & Shevach 2000),
3. Each Teff cell enhances the functions of those effector cells with which it shares a similar antigen specificity (Lanzavecchia 1985; Murphy et al. 2011), and
4. Each Treg cell suppresses effector cells found in its vicinity (i.e. bystander suppression) without regard to the particular HIV strains recognized by those cells (Shevach 2009; Thornton & Shevach 2000).

We argue that these experimental observations have the following important implication: the *relative* magnitude of the Teff response to HIV infection, compared to the Treg response, determines the effectiveness of HIV control by the immune system and, hence, the rate of AIDS progression. We define the magnitude of the Teff (Treg) response to HIV infection as the product of the concentration of responding Teff (Treg) cells and the efficacy of individual Teff (Treg) cells in promoting (suppressing) the elimination of HIV by the immune system. According to this theory, individuals whose anti-HIV immune responses are associated with a larger Teff:Treg response ratio will control HIV infection better and will consequently develop AIDS more slowly.

#### A mathematical instantiation of our hypothesis

We model effects of Treg and Teff cells on the rate at which an individual HIV patient progresses to AIDS. We quantify the AIDS progression rate using the rate of change of the total concentration of all HIV strains (i.e. the total number of HIV particles per ml of blood, also called the HIV load) found in a patient. We describe the dynamics of the concentration of the i^th^ HIV strain in the presence of immune responses mediated by both Treg and Teff cells by using the following system of ordinary differential equations

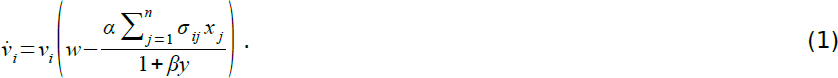

*v_i_* is the concentration of HIV strain *i*, *v̇_i_* is the time derivative representing the rate at which *v_i_* is changing, *x_i_* (*y_i_*) is the concentration of activated effector Teff (Treg) cells with specificity for strain *i*, *y* is the total concentration of all activated Treg cells, σ_*ij*_ is the probability that a T cell of “clonality” *j* will recognize HIV strain i, *n* is the number of different HIV strains, *α* is the efficacy of Teff cells in mediating HIV elimination, and *β* is the efficacy of Treg cells in suppressing immune responses. In the absence of specific immune responses the HIV load grows exponentially at the rate *w*. Teff cells promote effector responses in an HIV strain-specific manner, whereas Treg cells suppress those responses in a non strain-specific manner, as discussed earlier.

Summing Equation 1 over all possible HIV strains *i*, *i*=1,2,…,*n*, we obtain the following equation for the dynamics of the HIV load

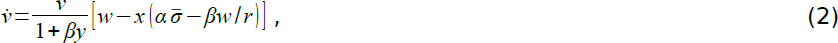

where

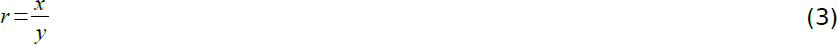

and

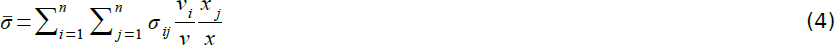

is a frequency-weighted average of the cross-reactivity that exists between pairs of HIV strains.

Equation 2 predicts that there are two possible patterns of AIDS progression, namely:

1. Progression to AIDS immediately after HIV infection. If *r* ≤ *βw/α*, then equation 2 predicts that the HIV load will not reach a steady-state value. Rather, the immune system will be unable to control HIV replication, allowing the HIV load to grow without bound soon after infection. We consider this state of inadequate control of HIV replication by the immune system to be analogous to AIDS. Therefore, the model predicts that a patient whose anti-HIV immune responses are associated with an r value that is smaller than or equal to one will develop AIDS soon after being infected.
2. Progression to AIDS after an incubation period. If *r > βw/α*, then equation 2 predicts that the HIV load will be regulated to a steady-state value. The Treg cell frequency will increase during the course of the infection (Montes et al. 2006; Mozos et al. 2007; Suchard et al. 2010), implying that r will decrease. Similarly, σ̄ will change during the infection, decreasing when a new HIV quasi-species emerges, increasing if the quasi-species takes over the entire HIV population, and then decreasing again during periods of further antigenic diversification. The rate of change of σ̄ will depend on the antigenic structure of HIV (e.g. it would be slow if on average mutations change only slightly the antigenic difference between the parent HIV strain and the mutant) as well as the strength of the immune response (e.g. it would be slow if the immune response is weak). A decrease in r, combined with a possible decrease in σ̄, would eventually cause the HIV load to increase uncontrollably, resulting in AIDS. This will occur when

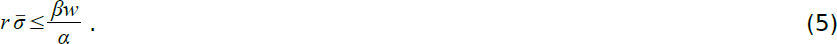 Therefore, the mathematical model predicts that a chronically infected HIV patient will develop AIDS when the product of the ratio of Teff vs. Treg cell frequencies and the average antigenic diversity becomes smaller than or equal to a threshold value that is proportional to both the Treg efficacy and the average HIV replication rate, but inversely proportional to the Teff efficacy.

The preceding analysis did not specify the dynamics of *x*, *y*, and other relevant variables. We confirm this simple analysis by developing a more detailed mathematical model (Methods). In addition to *v_i_*, the more detailed model also specifies the dynamics of 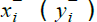, the concentration of resting memory Teff (Treg) cells of clonality *i*, and *x_i_* (*y_i_*), the concentration of activated Teff (Treg) cells of clonality i, (Methods).

A steady-state analysis of the more detailed model (Methods) yields the following threshold condition for the development of AIDS in individual patients

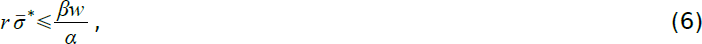

where σ̄* is the value of σ̄ evaluated at steady-state. Therefore, the steady-state behavior of the more detailed model predicts the same patterns of AIDS progression as described earlier.

We illustrate these patterns of AIDS progression by using numerical simulations of the more detailed model (Fig. 1). We simulated the dynamics of a single HIV strain (i.e. σ̄ = 1) infecting two patients, one (A) for whom the value of *r* is less than *βw/α* at the start of the simulation, and another (B) for whom *r* is greater than βw/α. As expected, patient A progresses to AIDS soon after infection (Fig. 1, left panel). In contrast, patient B progresses to AIDS only after a relatively long HIV incubation period (Fig. 1, left panel). In patient B, *r* decreases during infection (Fig. S1) because Treg cells are less susceptible to HIV-mediated depletion vs. Teff cells in this patient. Eventually, *r* becomes less than *βw/α*, leading to AIDS. To contrast the patterns just described with those expected in elite controllers, we show also results for a third patient (C) for whom *r* stays above *βw/α* throughout the infection (Fig. 1, left panel). The only parameter difference between patients B and C is that the Treg cells of B are five times less susceptible to HIV-mediated killing vs. the Treg cells of C. Notably, the total concentration of T cells that ever become activated is lowest in patient C (Fig. 1, right panel). These results illuminate the virus and immunologic parameters that produce different empirical patterns of AIDS progression within the framework of our hypothesis.

**Figure 1.**
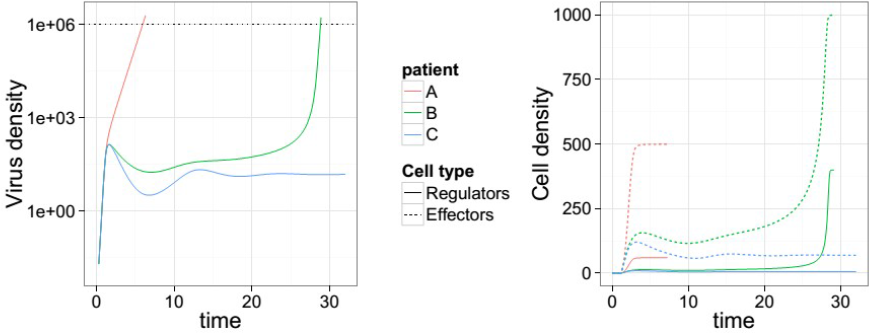
Mathematical model reproduces known empirical patterns of AIDS progression. We used the model given by Equations 1,7–10 to simulate HIV dynamics in three patients: A rapid AIDS progressor (A), a slower progressor (B), and a nonprogressor (C). To focus attention on the dependence of AIDS progression on r (as opposed to virus diversity), we simulated the dynamics of only one non-mutating HIV strain. AIDS progression is governed by both *r*_0_= *b*_x_/*b*_y_ and *k*_y_/*k*_x_, because *r* initially depends on *r*_0_, and it eventually converges to *r*_0_\**k*_y_/*k*_x_ when the HIV load becomes large [see Equations 11 & 12]. Therefore, we fixed the following parameters of the model: *α*=*β*=*b*_x_=*c*=λ=1, *f*=*k*_x_=*μ*_r_=0.001, *μ*=0.2, *w*=10. Then, we varied *b*_y_ and *k*_y_, as follows: Patient A: *b*_y_=0.12, *k*_y_=0.001; Patient B: *b*_y_=0.08, *k*_y_=0.0002; Patient C: *b*_y_=0.08, *k*_y_=0.001. We started each simulation with *x*=*x*^−^=*y*^−^=*y*=0, *v*=0.001. **Left panel:** Dynamics of HIV concentration in patients A-C. Initially *r*≈1/0.12 < 10 in patient A, leading to AIDS (i.e. uncontrolled virus replication) soon after infection. But, *r*≈1/0.08 > 10 initially in patients B & C, so AIDS does not occur. Meanwhile, as the HIV load grows r converges to ≈1/0.4 < 10 in patient B (Fig. S1), causing AIDS, but it does not change in patient C. **Right panel:** Dynamics of total concentrations of memory and activated Teff (dotted line) and Treg (solid line) cells found in patients A-C. Notably, the concentration of T cells that are ever activated by HIV is consistently lower in patient C vs. patients A & B.

#### Discussion

An improved understanding of the mechanisms that determine observed variation in the rates at which different untreated HIV patients develop AIDS will inform strategies for managing those patients. We described a hypothesis that sheds new light on the role of CD4^+^ T cells in the progression to AIDS. The basic tenet of our hypothesis is as follows: 1) Inadequate control of HIV replication causes AIDS; 2) CD4^+^ CD25^lo^ Foxp3^−^ T effector (Teff) cells enhance the functions of other effector immune cells (notably B and CD8^+^ T cells) that control HIV replication depending on the HIV strains recognized by those other cells (Lanzavecchia 1985; Murphy et al. 2011; Shedlock & Shen 2003); and 3) CD4^+^ CD25^hi^ Foxp3^+^ T regulatory (Treg) cells suppress the functions of the effector cells independent of the HIV strains that they recognize (Murphy et al. 2011; Shevach 2009; Thornton & Shevach 2000). The hypothesis proposes that individual variability in the ratio (r) of the Teff vs. Treg cell frequency causes variability in AIDS progression rates.

We used a mathematical instantiation of our hypothesis to define more precisely the relationship between r and the rate of AIDS progression. The mathematical model predicts that there are two possible modes of AIDS progression: On the one hand, the model predicts that a patient will develop AIDS soon after being infected with HIV if *r* σ̄ ≤ *βw/α*, where α is the efficacy of Teff cells in enhancing HIV elimination, *β* is the efficacy of Treg cells in suppressing HIV elimination, σ̄ is the probability that any two HIV strains are cross-reacting, and w is the average rate of HIV replication. A very fast progression to AIDS has previously been observed in some human HIV patients (CASCADE 2000). On the other hand, the model predicts that if *r* σ̄ is greater than *βw/α* at the time of HIV infection, then a patient may still develop AIDS if it eventually falls below *βw/α*. This might occur when *r* decreases during the course of infection (Mozos et al. 2007; Suchard et al. 2010; Nilsson et al. 2006) and σ̄ also decreases due to antigenic diversification of the HIV population found in a patient. The existence of this threshold is a generic consequence of our assumption that the enhancement of anti-HIV effector immune responses by Teff cells depends on the antigenic specificity of those responses, whereas the suppression of the same responses occurs in a “bystander” manner (Lanzavecchia 1985; Murphy et al. 2011; Shedlock & Shen 2003; Shevach 2009; Thornton & Shevach 2000). It does not depend on detailed assumptions. These theoretical results suggest that both HIV antigenic diversity (Nowak et al. 1990) and the ratio of Teff vs. Treg cell frequencies are important determinants of the AIDS progression rate.

These theoretical results are supported by our numerical simulations, which show (Fig. 1, left panel) that these different outcomes can result from variation in only two effective parameters: 1) the ratio of the frequency of naïve HIV-specific T conventional cells vs. the frequency of naïve HIV-specific Treg cells (*b*_x_/*b*_y_); and 2) the ratio of the rate of HIV-mediated killing of Treg vs. Teff cells (*k*_y_/*k*_x_). The value of *r* depends on *r*_o_ = *b*_x_/*b*_y_ at the time of infection, and on r_o_*k_y_/k_x_ at later time points when the HIV load becomes large. The simulations highlight, in particular, the key roles that naïve T-cell repertoire variability (Moon et al. 2007) and differential T-cell susceptibility to HIV-mediated depletion might play in generating different clinical outcomes of HIV infection.

Our hypothesis provides plausible mechanistic explanations for various other empirical facts concerning variability in AIDS progression rates. Three examples are discussed below.

First, our hypothesis provides an explanation for the observation that in the absence of treatment older adults who are immunocompetent develop AIDS after HIV infection at a significantly faster rate than immunocompetent younger adults (CASCADE 2000; Van der Paal et al. 2007). Past studies show that the frequency of blood-circulating Treg cells increases in immunocompetent adults during aging (Gregg et al. 2005; Lages et al. 2008; Trzonkowski et al. 2006). HIV infection promotes the recruitment of such circulating Treg cells into lymphoid tissues (Andersson et al. 2005), where they are able to suppress the development of anti-HIV effector immune responses. This implies that the anti-HIV immune responses of older adults may be associated with smaller values of *r* compared to the responses mounted by younger adults. Therefore, according to our hypothesis, untreated older adult HIV patients will develop AIDS faster than untreated younger adult patients.

Second, our hypothesis provides an explanation for the observation that the infection of an HIV-infected individual by certain other pathogens, including other HIV variants, accelerates the development of AIDS (Cornelissen et al. 2012; Toossi et al. 2001). Specifically, other pathogens that infect similar tissues as HIV will induce Treg cells that suppress not only the Teff responses to those pathogens, but also the Teff responses to HIV. In contrast, if those pathogens are antigenically dissimilar to the resident HIV strain, then their induced Teff responses will be ineffective against that strain. The resulting increase in the Treg cell concentration without a corresponding increase in the Teff cell concentration will decrease the *r* value associated with anti-HIV immune responses. According to our hypothesis, this decrease in *r* will accelerate the development of AIDS in the dually infected patient.

Third, our hypothesis suggests that natural hosts of simian immunodeficiency virus (SIV) have a higher *r* value compared to non-natural hosts that develop AIDS. For example, SIV infection is asymptomatic in African green monkeys, its natural host, but pathogenic in pigtailed macaques (Favre et al. 2009). Consistent with our hypothesis, experimental data indicate that SIV infection in the macaques results in a dramatic decrease in the ratio of the frequency of Th17 effector T cells vs. Treg cells, but this does not occur during infection of the green monkeys (Favre et al. 2009). Thus, our hypothesis provides a parsimonious explanation for the observed differential pathogenicity of SIV infection in the considered examples of natural vs. non-natural hosts.

Our hypothesis has implications for the therapeutic management of HIV patients. First, it predicts that AIDS might be delayed by increasing the ratio of effector to regulatory T cells, r. This can be increased by depleting HIV-specific Treg cells or augmenting HIV-specific Teff cells. Non-specific depletion of Treg cells has already proven useful in significantly reducing viral loads in animal models of retrovirus infections (Dietze et al. 2011; Zelinskyy & Dietze 2009), and also in improving effector immune responses to cancer in human patients (Mahnke et al. 2007). A possible strategy for specific deletion of HIV-specific Treg cells could be through the use of antibodies targeting the dominant Vbeta segment found in the Treg antigen receptors (Zaller et al. 1990). In addition, the hypothesis predicts that AIDS might be delayed by reducing *βw/α* – e.g., by using anti-CD39 antibodies to deplete the CD39 molecule that is expressed by Treg cells and whose amount is correlated with *β* (Deaglio et al. 2007; zur Wiesch et al. 2011). Furthermore, measuring the threshold value defined in Equation 5 might allow individual patients to be classified according to their projected risk for developing AIDS in order to better tailor therapy to individuals. The parameters of the threshold value can be approximated using in vitro experiments; *w* as the average replication rate of HIV found in a patient’s blood sample (Wolinsky et al. 1996); *α* as the rate at which a standardized or fixed amount of a patient’s Teff cells enhance the killing rate of infected cells by a fixed amount of naïve HIV-specific CD8+ T cells; and *β* as the factor by which a fixed amount of the same patient’s Treg cells reduces that cell killing rate.

In summary, we introduced a hypothesis that provides a mechanistic explanation for different empirical patterns of AIDS progression. It predicts that an HIV patient will develop AIDS if *r* is less than a defined threshold, implying that individual variation in *r* causes observed variation in AIDS progression rates. In addition, we identified certain alterable molecular factors that are mechanistically associated with AIDS progression in individual patients, including: 1) The frequency of the predominant T-cell receptor Vbeta gene that is found in a patient’s HIV-specific Treg cells, and 2) the level of CD39 expression by a patient’s Treg cells. Altering one or more of these molecular factors in individual patients so that *r* stays above the threshold represents a potential strategy for delaying AIDS. These predictions of our hypothesis warrant further theoretical, empirical, and experimental investigation.

## Methods

### A more detailed mathematical instantiation of our hypothesis

In addition to Equation (1), the model consists of the following equations

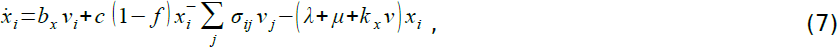

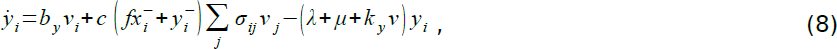

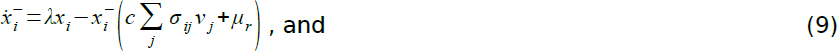

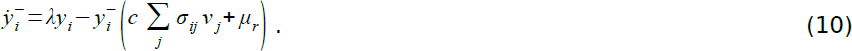

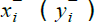 is the concentration of resting memory Teff (Treg) cells of clonality *i*, *x_i_* (*y_i_*) is the concentration of activated Teff (Treg) cells of clonality *i*, *i*=1,2,…,*n, μ* (*μ_r_*) is the natural death rate of activated (resting memory) T cells, *λ* is the rate at which activated T cells return to the resting, memory state, and *k_x_v* (*k_y_v*) is the rate at which activated Teff (Treg) cells are killed by HIV. Teff (Treg) cells of clonality *i* are activated from naïve conventional T cells (naïve Treg cells) at the rate *b_x_v_i_* (*b_y_v_i_*), and from resting memory Teff (Treg) cells at the rate 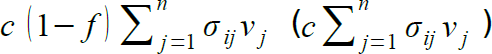. A certain fraction f of memory Teff cells are converted into Treg cells during the course of their activation (Vukmanovic-Stejic & Zhang 2006). Naïve T cell pools of different clonalities are assumed to be of the same size and constant over time.

We analyze the steady-state behavior of the mathematical model, assuming that resting memory T cells die at a much slower rate than they are activated by HIV strains – i.e. we assume that *μ_r_* is a slow variable and set *μr* = 0. The steady-state concentrations of activated Teff and Treg cells of clonality i are respectively

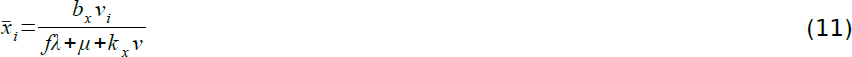

and

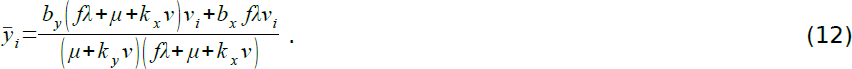

Substituting *x*̄*_i_* and *y*̄*_i_*into Equation 1 and summing over all possible HIV strains *i*, *i=*1,..,*n*, leads to the following equation for the dynamics of the steady-state HIV load

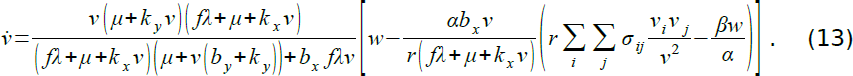

Note that at steady-state, *x_i_ /X* = *v_j_*/*v*, implying that

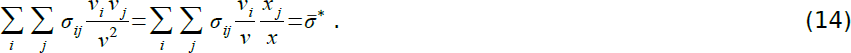

## Acknowledgments

The work described here was partially funded by a postdoctoral research fellowship from the German Academic Exchange Service (to WN). WN occupies a Joint Career Development Chair at AIMS Centres in South Africa and Ghana, funded by the Canadian International Development Research Centre. JD holds a New Investigator award from the Canadian Institutes for Health Research.

